# Selective reorganization of face-processing lateralization in participants with right-lateralized language network

**DOI:** 10.64898/2026.02.17.706260

**Authors:** Parham Zarger, Reza Rajimehr

## Abstract

A well-known feature of cortical organization is the lateralization of the language network (LN) to the left hemisphere (LH). Although atypical LN lateralization to the right hemisphere (RH) is well documented, its consequences for other lateralized functions remain poorly understood. Face processing and other category-selective visual areas, which are typically RH-biased, provide key test cases for examining whether atypical LN lateralization is associated with reorganization in other networks. Using functional magnetic resonance imaging (fMRI) data from the Human Connectome Project (HCP), we identified participants with right-hemisphere language dominance (RHLD) and examined whether category-selective visual areas exhibit reorganized hemispheric biases in these participants relative to those with typical left-hemisphere language dominance (LHLD). The results demonstrated selective reorganization within the face-selective areas: some regions showed a leftward shift in hemispheric bias in RHLD participants, whereas others preserved the canonical RH preference. This pattern indicates that cortical reorganization associated with atypical LN lateralization is region-specific rather than global, consistent with a flexible neural architecture in which hemispheric specialization can be selectively reconfigured. These results clarify how atypical LN lateralization impacts the hemispheric organization of face-selective areas and provide evidence for altered network-level specialization in the cerebral cortex.

## Introduction

The human language network (LN) comprises a set of brain regions that collectively support the core components of language processing, including comprehension, production, and integration of linguistic information (Binder et al., 1997; Fedorenko et al., 2011). The LN is primarily distributed across lateral temporal and lateral frontal areas and shows robust left hemisphere (LH) lateralization in most participants (Friederici, 2011). Despite inter-individual variability in functional topography, this hemispheric asymmetry is a highly consistent feature of human brain organization (Fedorenko et al., 2024). However, a minority of participants exhibit atypical LN lateralization, with right hemisphere (RH) dominance or bilateral distribution. The factors underlying these departures from the canonical pattern remain incompletely understood (Ocklenburg et al., 2013; Sha et al., 2021).

The reversal of LN lateralization in such participants raises fundamental questions about whether this atypical hemispheric organization extends beyond language to other lateralized cognitive functions. Specifically, if LN lateralization is altered, do other lateralized functions undergo a corresponding global reversal, exhibit selective reconfiguration, or remain unaffected? Addressing these questions is critical for determining whether atypical LN lateralization is associated with coordinated reorganization across functionally distinct networks or instead reflects region-specific reconfiguration.

To investigate the functional consequences of atypical LN lateralization, we focused on category-selective visual areas, which offer a well-characterized system for examining hemispheric asymmetries beyond language. These areas show reliable lateralization across participants. For instance, face-selective areas in the fusiform gyrus (fusiform face area, FFA) and superior temporal sulcus (STS) are typically right-lateralized (Kanwisher et al., 1997; Pitcher et al., 2011; Yovel et al., 2008). Similarly, other areas, such as place– and body-selective regions, tend to show a comparable hemispheric bias. The parahippocampal place area (PPA) which is involved in recognizing spatial layouts and places, and the extrastriate body area (EBA) which is involved in body recognition, both exhibit this canonical pattern (Downing et al., 2001; Epstein and Kanwisher, 1998). These cortical systems therefore provide an opportunity to assess whether the atypical LN lateralization is associated with coordinated or selective changes in the hemispheric organization of non-language functions.

Determining whether reorganization is coordinated or selective requires direct measures of lateralization for both LN and category-selective visual networks within the same participants. Despite extensive research on hemispheric asymmetries, evidence directly linking the lateralization of these two networks remains surprisingly limited. Previous works have mostly examined associations between handedness and the lateralization of some category-selective visual areas (Bukowski et al., 2013; Willems et al., 2010). Although handedness has historically been linked to LN lateralization—with left-handed participants showing higher rates of atypical RH language dominance (Knecht, 2000)—recent findings indicate that handedness may not reliably predict LN lateralization (Packheiser et al., 2020). This leaves a critical gap: whether hemispheric biases in category-selective visual areas covary with LN lateralization when the latter is measured directly through functional neuroimaging. Furthermore, no study has systematically examined whether atypical LN lateralization is associated with coordinated reorganization across multiple category-selective visual areas or whether such reorganization occurs selectively in specific regions. Here, we addressed these questions by identifying right-hemisphere language dominance (RHLD) participants using functional magnetic resonance imaging (fMRI) measures of LN lateralization and testing hemispheric biases in face-, place-, and body-selective areas. We hypothesized that if cortical reorganization associated with atypical LN lateralization operates globally, all category-selective visual areas in RHLD participants should show a leftward shift. Alternatively, if reorganization is region-specific, some areas may shift toward the LH while others preserve the canonical RH preference, revealing a more flexible and differentiated pattern of neural reorganization.

## Results

To test whether the reversal of LN lateralization is accompanied by changes in category-selective visual areas, we analyzed fMRI data from the Human Connectome Project (HCP), a large-scale publicly available dataset mapping structure, function, and connectivity of the human brain (Van Essen et al., 2013). The cortical maps were analyzed on a standard surface where LH and RH were registered to each other (Glasser et al., 2016).

We began by localizing the LN using the group-average activation map from the story vs. baseline contrast in the language processing task (Figure 1A). This task involved participants listening to brief auditory stories and then answering a two-alternative forced-choice comprehension question about the story content (Binder et al., 2011).

**Figure 1.**
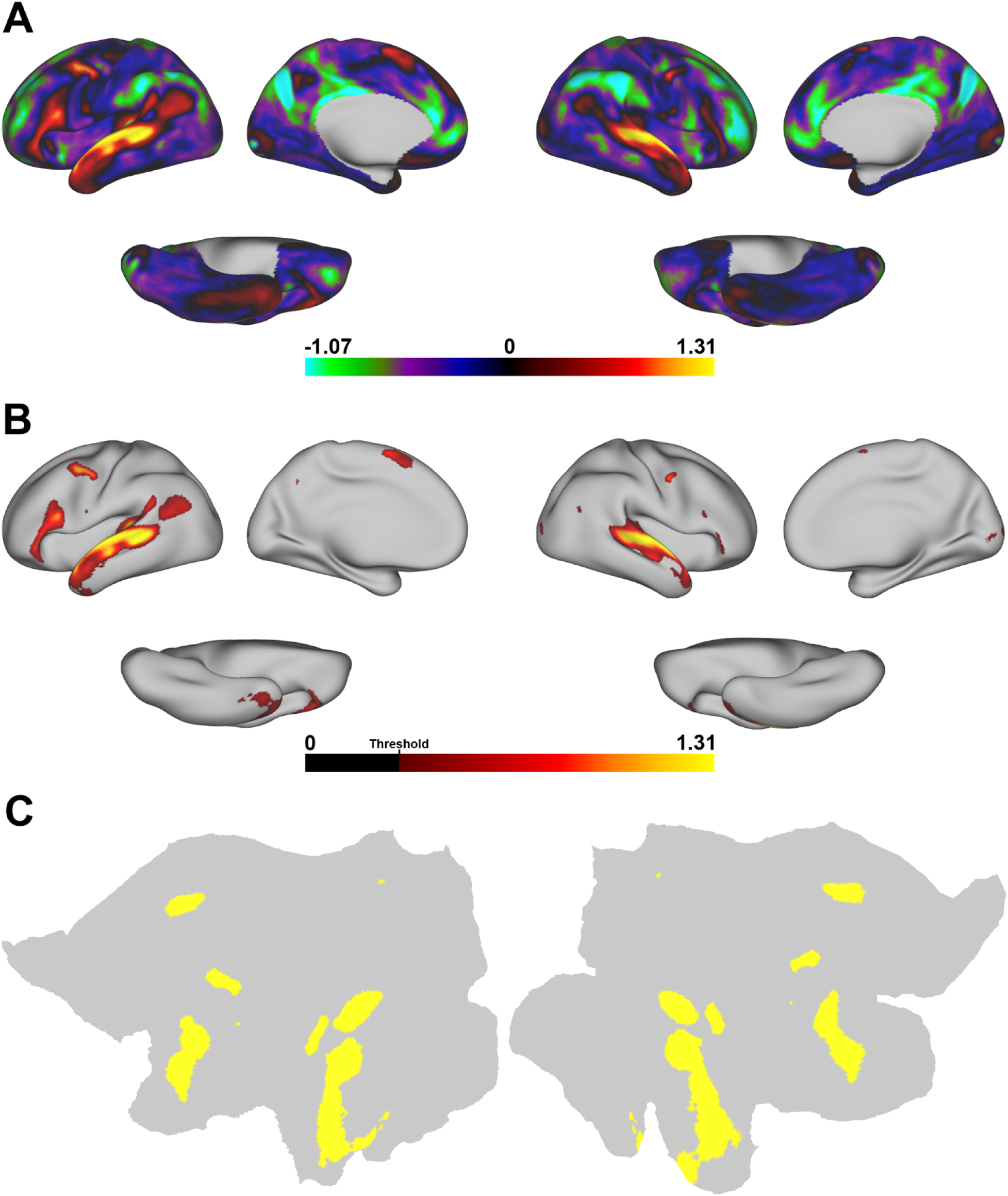
Bilateral language mask used in the laterality analysis. (A) Group-average activation map displaying the contrast between the story versus baseline conditions from the HCP language task. The map shows Cohen’s d effect sizes, displayed on the lateral, medial, and ventral views of the cortical surface for both hemispheres. In each view, the LH is shown on the left, and the RH is shown on the right. The color scale bar ranges from negative effect sizes (cyan/blue) to positive effect sizes (red/yellow), reflecting regions of decreased or increased activation, respectively. (B) To localize language-responsive areas, a threshold of Cohen’s d > 0.3 was applied to the activation map. This threshold highlights regions with small to medium effect sizes, ensuring the inclusion of all language-responsive vertices. As in Panel A, the activation map is displayed on lateral, medial, and ventral views of the cortical surface. The color scale bar ranges from 0.3 to 1.31, indicating regions of increased activation above the threshold, aimed at delineating the full extent of the language network. (C) A two-dimensional (2D) flat projection of the LH and RH showing the final language mask. This mask was created by applying a threshold of Cohen’s d > 0.3 to the group-average activation map for the story versus baseline contrast, excluding early sensory regions such as the auditory and visual areas. The language-specific vertices from both hemispheres were then combined to form a bilateral mask (i.e., identical/homotopic regions in LH and RH). This mask was subsequently used to calculate the LLI.

To delineate the extent of language-responsive network, we applied a threshold of Cohen’s d > 0.3 to the group-average activation map (Figure 1B). This threshold was used to generate an initial map of task-responsive regions, which was subsequently refined to derive a more specific definition of the LN. As expected, given the use of spoken narrative stimuli, the thresholded map included activations within early sensory areas, most notably along the auditory cortex, with comparatively sparse activation in the early visual cortex. The auditory responses likely reflect obligatory acoustic and speech-sound processing evoked by the stories rather than language-selective computations *per se* (Hickok and Poeppel, 2007; Price, 2012). The weaker visual responses may reflect top-down influences such as visual imagery or attentional engagement elicited by the narratives (Kosslyn et al., 2001). Consistent with this interpretation, recent works emphasize a functional distinction between the LN and early sensory areas. In spoken story comprehension, early auditory regions are expected to respond strongly because they encode acoustic and speech-sound features, whereas the LN supports higher-level linguistic computations operating over these inputs (Fedorenko et al., 2024). Therefore, including early sensory regions in our definition would increase the risk of mixing sensory-driven activity with LN responses, potentially obscuring the functional topography of the LN (Fedorenko et al., 2011). To avoid conflating sensory-driven responses with LN activity, we excluded early sensory regions from the thresholded activation map using the multimodal cortical parcellation (Glasser et al., 2016), thereby focusing the analysis on the core LN (Binder, 2000; Friederici, 2011). Given the one-to-one correspondence between LH and RH vertices, the remaining language-responsive vertices were combined across both hemispheres to form a single bilateral LN mask (Figure 1C). This unified mask served as the basis for calculating the language laterality index (LLI) for each participant in subsequent analyses, ensuring a consistent and anatomically matched definition of the language network across hemispheres and participants.

Next, we examined hemispheric lateralization within the core LN across all participants (N = 1049) by computing the LLI. For each participant, activation values were extracted separately from the LH and RH portions of the predefined bilateral LN mask. The LLI was then calculated as the difference between mean activation in the LH and mean activation in the RH, with positive values indicating LH dominance and negative values indicating RH dominance.

Based on the distribution of LLI values across all participants (Figure 2A), we defined two groups exhibiting strong hemispheric dominance for language: left-hemisphere language dominance (LHLD) and right-hemisphere language dominance (RHLD). Participants with LLI values more than two standard deviations above or below the mean were classified as LHLD or RHLD, respectively. This classification yielded 17 LHLD participants and 28 RHLD participants. To enable a balanced comparison between groups, we supplemented the LHLD group with 11 additional participants exhibiting the highest positive LLI values, resulting in 28 participants per each group. This procedure restricted the analysis to participants with the strongest LN lateralization while maintaining matched group sizes for subsequent comparisons.

**Figure 2.**
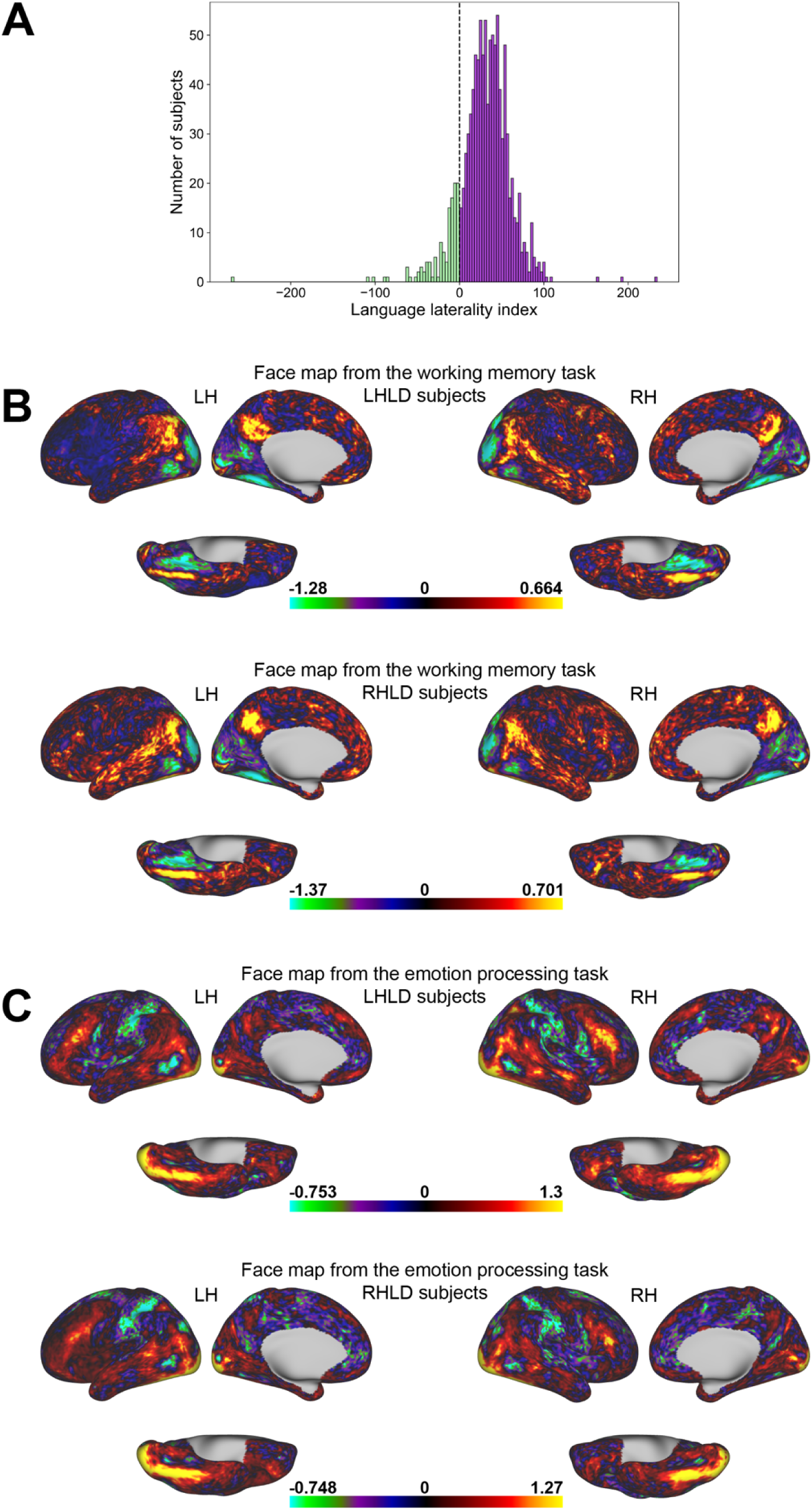
Language laterality index across all participants and group-average face maps in LHLD and RHLD participants. (A) Distribution of language laterality index (LLI) across all participants (N = 1049). The histogram illustrates the hemispheric dominance for language processing, with the LLI calculated by subtracting the average activation in the RH language mask from the LH language mask. Positive values indicate left-hemisphere language dominance, while negative values indicate right-hemisphere language dominance. Two subgroups were identified: left-hemisphere language dominance (LHLD) and right-hemisphere language dominance (RHLD) participants. The threshold for classification was set at two standard deviations from the mean LLI values, with participants beyond this threshold classified as strongly LHLD or RHLD. The histogram bin width was determined using the Freedman-Diaconis rule to ensure optimal representation of the data distribution. (B) Cohen’s d effect size maps for the working memory task (FACE-AVG contrast) in LHLD and RHLD participants. (C) Cohen’s d effect size maps for the emotion processing task (faces vs. shapes contrast) in LHLD and RHLD participants.

Building on the classification of LHLD and RHLD participants, we next examined whether differences in LN lateralization were associated with differences in hemispheric bias within the face-processing network. To this end, we analyzed brain activation patterns for faces during two relevant tasks from the HCP: the working memory task using the FACE-AVG contrast (faces vs. places, tools, and body parts) and the emotion processing task using the faces vs. shapes contrast. These tasks were selected because they reliably engage canonical face-selective areas and provide complementary probes of face processing under cognitive and affective demands. The working memory task consisted of blocks of images from four categories (faces, places, tools, and body parts), performed under 0-back and 2-back working memory conditions (Barch et al., 2013). The emotion processing task required participants to match either emotional faces (angry or fearful) or geometric shapes across block types (Hariri et al., 2006).

We then compared group-average activation patterns between LHLD and RHLD participants by generating Cohen’s d maps separately for each group. In LHLD participants, face-related activations in occipito-temporal cortex showed a clear RH bias (Figure 2B), whereas in RHLD participants, this activity appeared to be biased toward the LH (Figure 2C).

To characterize these hemispheric biases at a finer spatial scale, we conducted region-of-interest (ROI) analyses focusing on canonical face-selective areas. These areas were defined based on the group-average FACE-AVG contrast from the working memory task, applying a threshold of Cohen’s d > 0.3. The localized ROIs included the posterior superior temporal sulcus (pSTS), middle superior temporal sulcus (mSTS), anterior superior temporal sulcus (aSTS), medial face area (MFA), occipital face area (OFA), and fusiform face area (FFA). These regions are well-established components of the face-processing network (Allison et al., 2000; Grill-Spector and Malach, 2004; Haxby et al., 2000; Kanwisher et al., 1997; Pitcher et al., 2011). The ROIs were merged across both hemispheres to allow direct comparison of lateralization patterns between LHLD and RHLD participants (Figure 3A). After defining the face-selective ROIs, we quantified hemispheric bias by computing a face laterality index (FLI) for each participant. For each ROI, the FLI was calculated as the difference between mean activation in the LH and RH, using activation values extracted from the corresponding ROIs. This procedure was performed separately for the working memory and emotion processing tasks. Positive FLI values indicate greater LH activation, whereas negative FLI values indicate greater RH activation.

**Figure 3.**
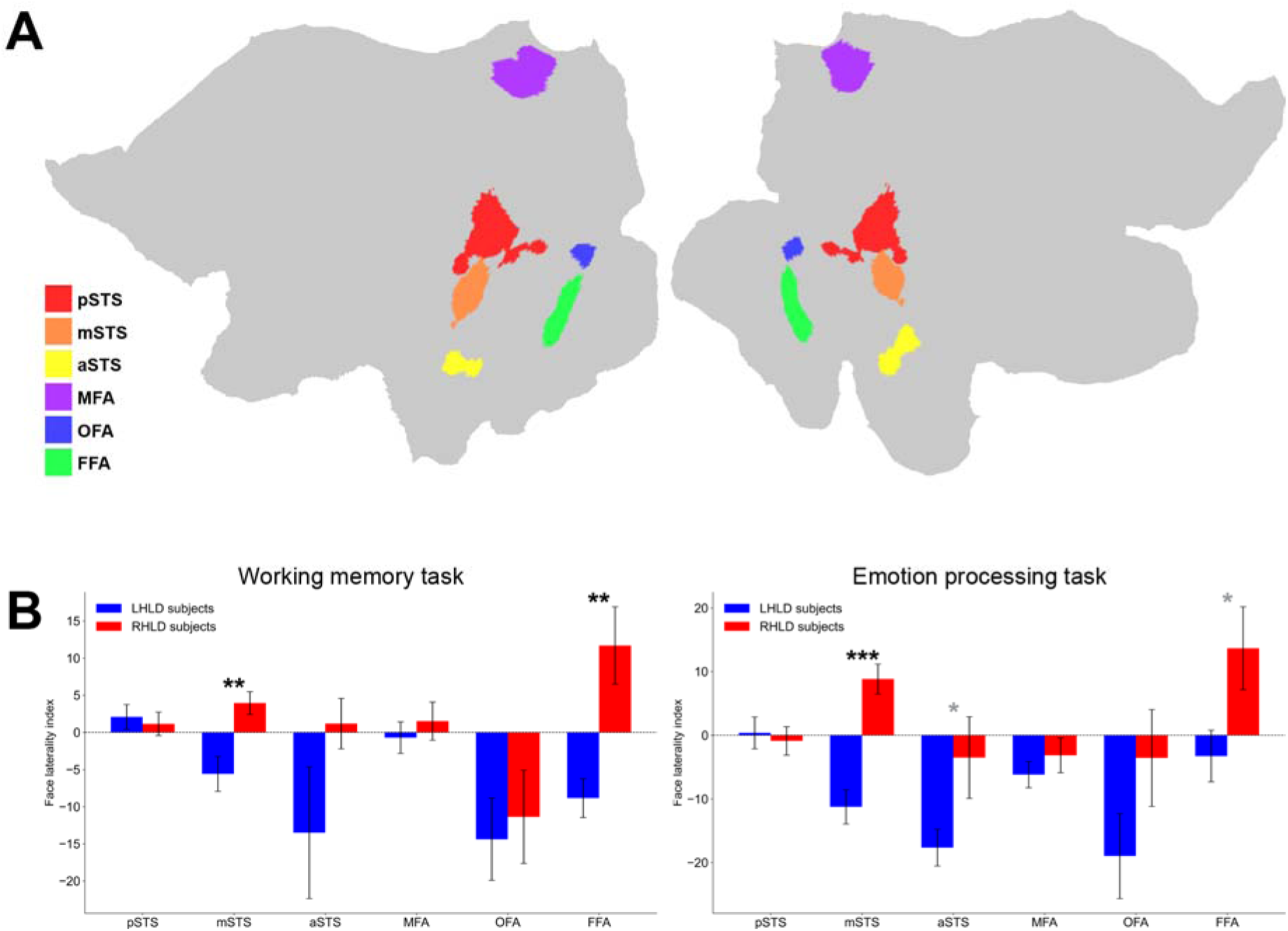
ROI analysis of face laterality index in LHLD and RHLD participants. (A) Flat patch visualization of the regions of interest (ROIs) used to extract activation values for each subject. The ROIs include the posterior superior temporal sulcus (pSTS, red), middle superior temporal sulcus (mSTS, orange), anterior superior temporal sulcus (aSTS, yellow), medial face area (MFA, purple), occipital face area (OFA, blue), and fusiform face area (FFA, green). (B) Group differences in the face laterality index (FLI) between LHLD (blue) and RHLD (red) participants across different ROIs during the working memory task (left) and the emotion processing task (right). The error bars indicate one standard error of the mean (SEM). Significant differences between LHLD and RHLD participants are indicated with asterisks (*p < 0.05, **p < 0.01, ***p < 0.001; black asterisks: FDR-corrected significance, gray asterisks: uncorrected significance).

Significant differences in FLI between LHLD and RHLD groups were observed in the mSTS (p = 0.001; FDR-corrected p = 0.003) and FFA (p = 0.001; FDR-corrected p = 0.003) during the working memory task. Face activations in the mSTS and FFA were RH-biased in LHLD participants and LH-biased in RHLD participants. No differences were detected in the pSTS (p = 0.686), aSTS (p = 0.128), MFA (p = 0.508), and OFA (p = 0.721). During the emotion processing task, the mSTS again showed a significant difference (p = 7 × 10□□; FDR-corrected p = 4 × 10□□). In contrast, the FFA did not remain significant after FDR correction (p = 0.031; FDR-corrected p = 0.093), and the aSTS showed a marginal effect prior to correction (p = 0.049) that did not survive FDR correction (FDR-corrected p = 0.098). No differences were observed in the pSTS (p = 0.708), MFA (p = 0.380), and OFA (p = 0.411) during the emotion processing task. Overall, pSTS, MFA, and OFA showed no evidence of differential hemispheric bias between LHLD and RHLD groups across either task. Across both tasks, the mSTS and FFA were the only regions showing a significant difference (Figure 3B).

To ensure that the observed effects were not driven by a specific activation threshold, we analyzed the FLI across varying percentages of the most active vertices in each bilateral ROI, ranging from 100% to 5%. This analysis was conducted for both the mSTS and FFA across the two tasks. The mSTS lateralization profile remained stable across all thresholds: RHLD participants consistently showed greater LH activation (positive FLI); LHLD participants consistently showed greater RH activation (negative FLI). In the mSTS, this profile persisted across the entire range of selected vertices in both tasks, indicating that the mSTS laterality effect is not dependent on the threshold used (Figure 4A). Similarly, in the FFA, the direction of hemispheric lateralization was consistent across all thresholds in both tasks. The preservation of this effect across activation thresholds indicates that the observed FFA bias is not dependent on a specific vertex-selection cutoff (Figure 4B). These findings demonstrate that hemispheric differences in mSTS and FFA are stable across vertex-selection thresholds.

**Figure 4.**
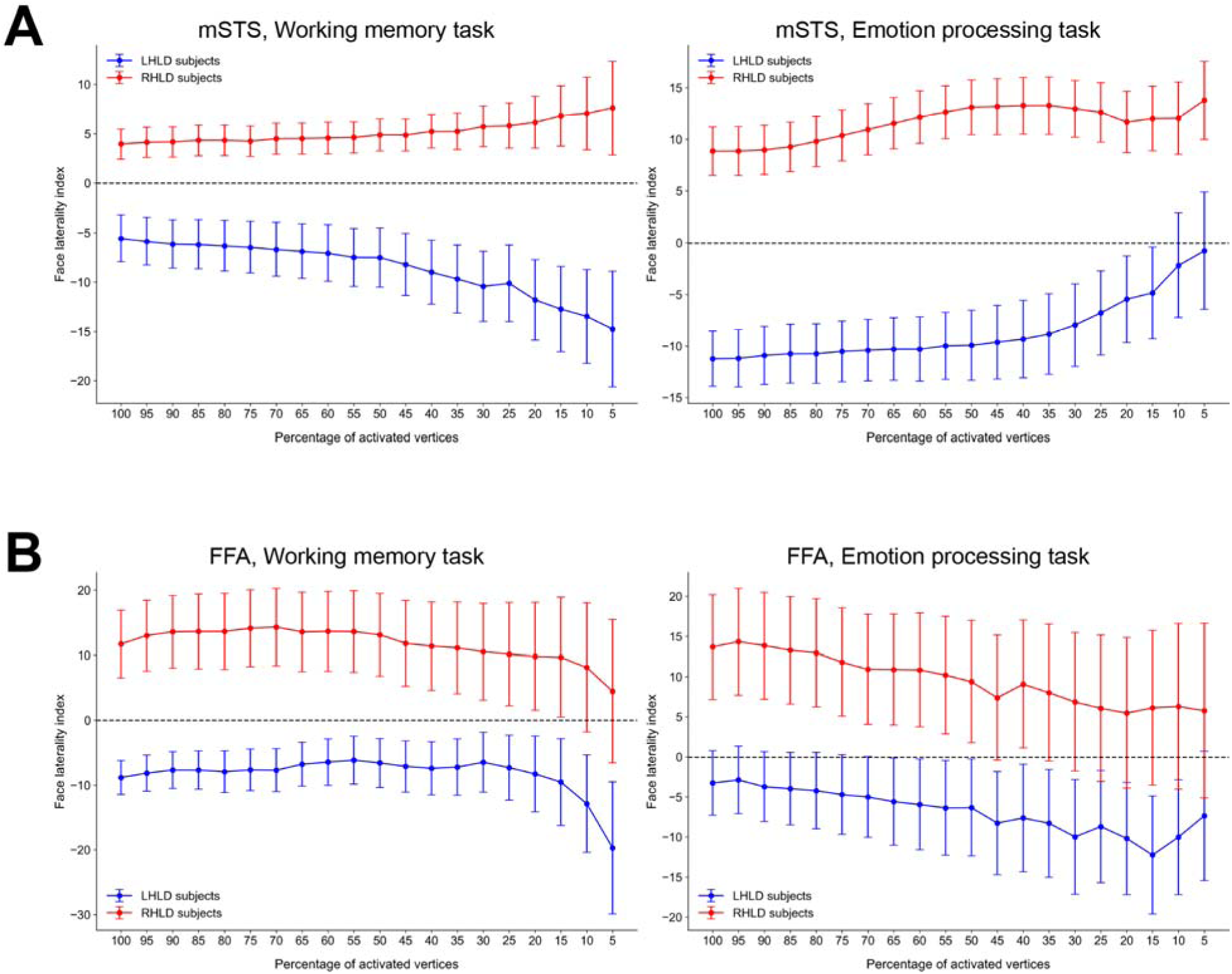
Robustness of the lateralization effects across varying levels of selected vertices. In each ROI, we systematically sampled face-selective vertices, ranging from 100% of vertices to 5% of most active vertices in the group-average face localizer map. For each group of subjects (LHLD and RHLD), we then plotted the FLI as a function of different percentages of activated vertices in the mSTS (A) and FFA (B) during the working memory task and emotion processing task. The left panel shows the FLI for the mSTS during the working memory task. The error bars indicate one standard error of the mean (SEM).

To determine whether group-level differences reflect a continuous relationship between language and face-processing network lateralization across all participants, we correlated each participant’s LLI with their FLI in the mSTS and FFA across both tasks (working memory: n = 1,036; emotion processing: n = 1,040). The results revealed negative correlations between LN and face-processing network lateralization in both regions. In the mSTS, greater LH language dominance was associated with greater RH face-processing bias in both the working memory task (r = −0.15, p < 0.001) and the emotion processing task (r = −0.19, p < 0.001) (Figure 5A). The FFA showed a parallel pattern, with negative correlations in both the working memory task (r = −0.14, p < 0.001) and the emotion processing task (r = −0.10, p < 0.01) (Figure 5B). These correlations indicate that the relationship between language and face-processing lateralization is not limited to categorical group differences between LHLD and RHLD participants, but instead reflects a graded, continuous relationship across the full spectrum of LN lateralization. The mSTS showed the most consistent association across tasks, whereas the FFA exhibited a reliable but weaker association, consistent with the group-level analyses.

**Figure 5.**
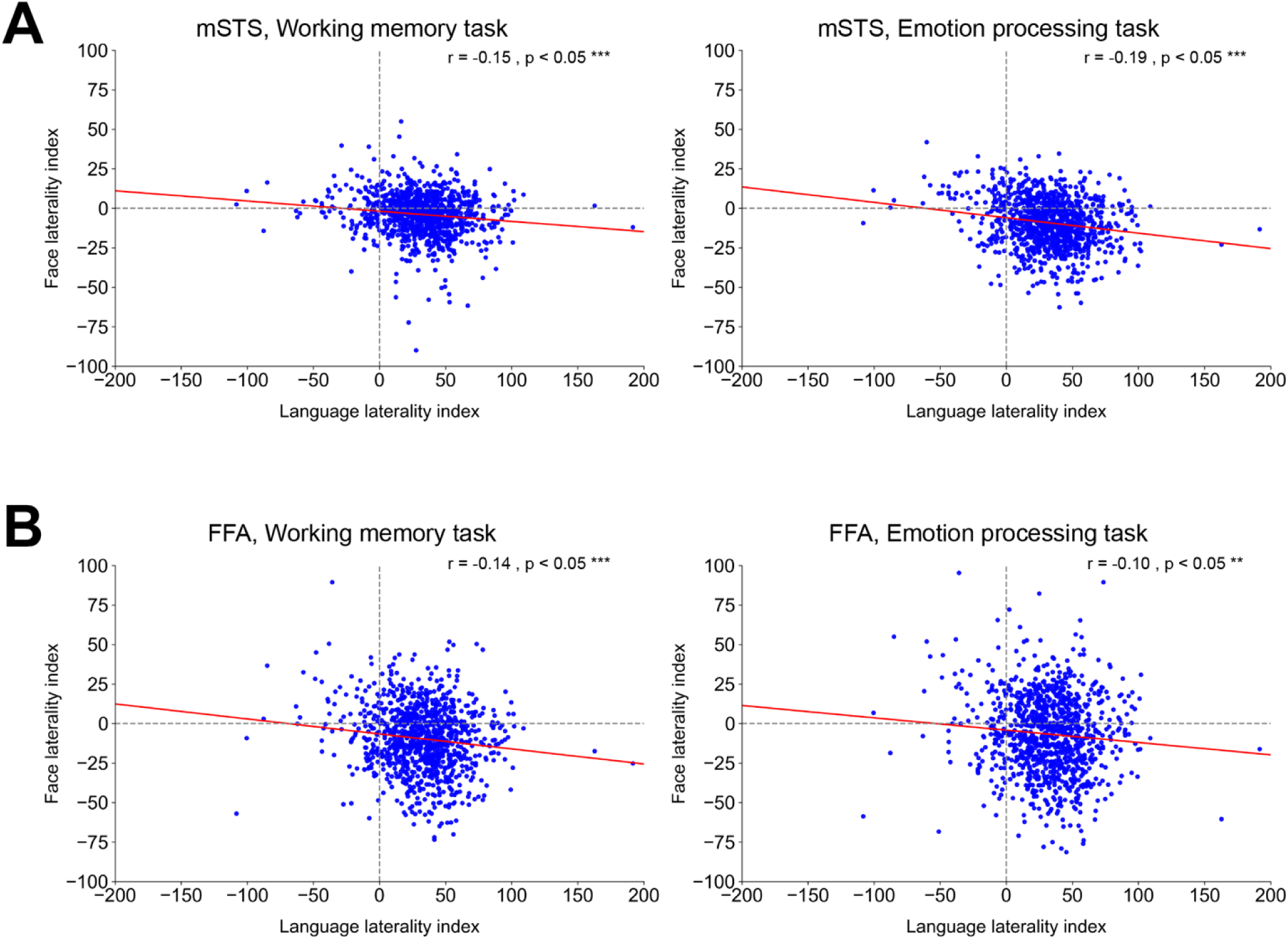
Relationship between language laterality index and face laterality index across all participants. (A) Scatterplots showing the relationship between the language laterality index (LLI) and the face laterality index (FLI) in the middle superior temporal sulcus (mSTS) across participants during working memory task (left) and emotion processing task (right). (B) Scatterplots showing the relationship between LLI and FLI in the fusiform face area (FFA) across participants during working memory task (left) and emotion processing task (right). The red lines are regression lines. Significant correlations are indicated with asterisks (*p < 0.05, **p < 0.01, ***p < 0.001).

Having demonstrated that LN lateralization is associated with systematic differences in hemispheric bias within face-selective areas, we next examined whether this relationship extends to other category-selective visual networks. Specifically, we tested whether place– and body-selective areas show comparable LHLD–RHLD differences in hemispheric bias, or whether such differences are restricted to face-processing network.

We identified place-selective and body-selective areas using the same approach applied to face-selective areas. For place-selective areas, we used the group-average PLACE-AVG contrast from the working memory task, applying a threshold of Cohen’s d > 0.6 to restrict the analysis to vertices exhibiting moderate-to-large effect sizes. To exclude low-level visual processing regions, vertices corresponding to early visual cortex (V1–V3) were removed, thereby isolating higher-level regions. This procedure identified canonical place-selective ROIs, including the occipital place area (OPA), parahippocampal place area (PPA), medial place area (MPA), and posterior intraparietal gyrus place area (PIGPA) (Epstein et al., 1999; Kamps et al., 2016; Kennedy et al., 2022) (Figure 6A). For body-selective areas, we applied the same Cohen’s d > 0.6 threshold to the group-average BODY-AVG contrast, which delineated the extrastriate body area (EBA) and fusiform body area (FBA) (Downing et al., 2001; Peelen and Downing, 2005) (Figure 6B). As with face-selective areas, ROIs were merged bilaterally to enable comparison of lateralization patterns between LHLD and RHLD participants.

**Figure 6.**
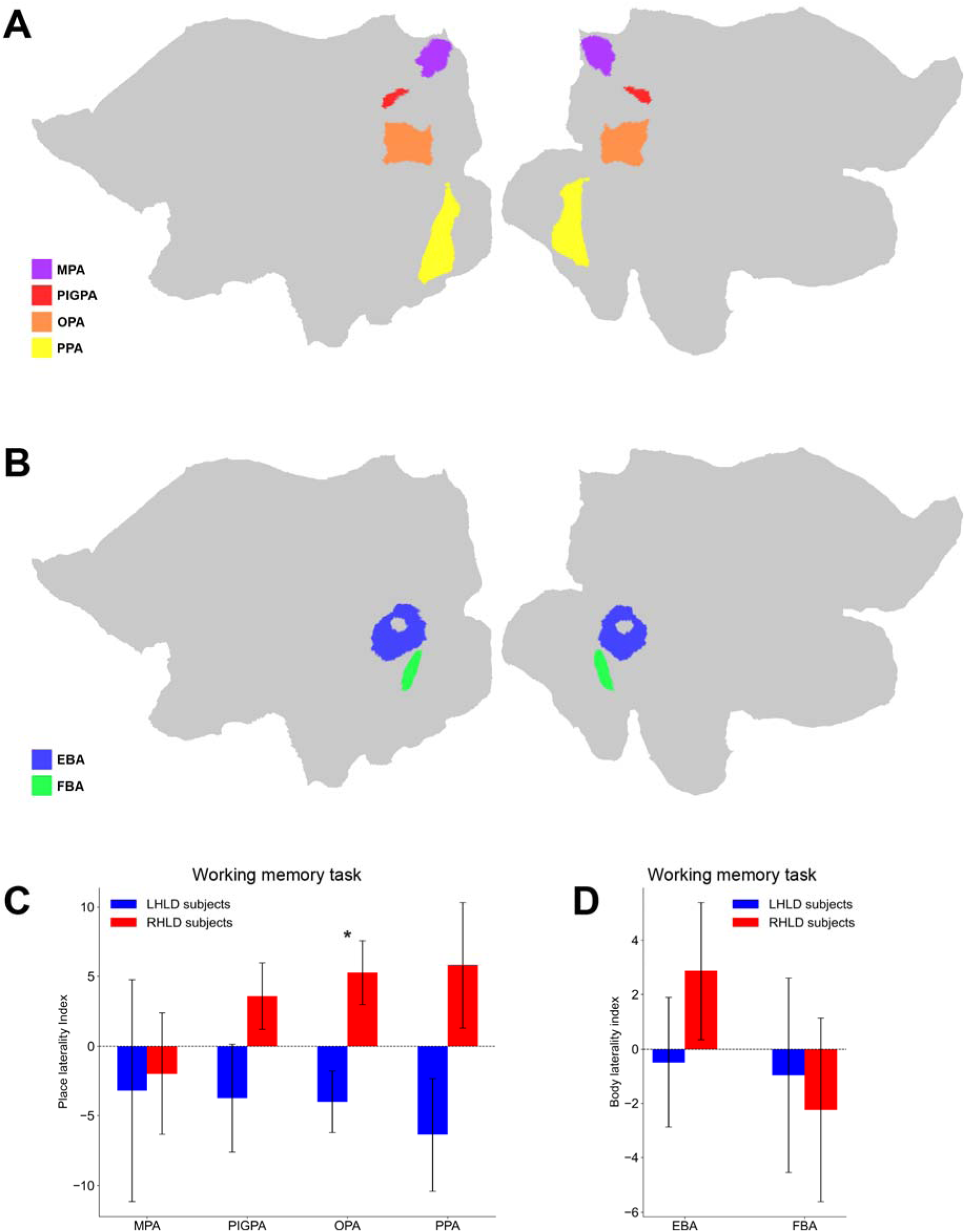
ROI analysis of place and body laterality index in LHLD and RHLD participants. (A) Flat patch visualization of the place-selective regions of interest (ROIs) used to extract activation values for each subject. The ROIs include the medial place area (MPA, purple), posterior intraparietal gyrus place area (PIGPA, red), occipital place area (OPA, orange), and parahippocampal place area (PPA, yellow). (B) Flat patch visualization of the body-selective ROIs used to extract activation values for each subject. The ROIs include the extrastriate body area (EBA, blue) and fusiform body area (FBA, green). (C–D) Group differences in the place laterality index (C) and body laterality index (D) between LHLD (blue) and RHLD (red) participants across different ROIs during the working memory task. The error bars indicate one standard error of the mean (SEM). Significant differences between LHLD and RHLD participants are indicated with asterisks (*p < 0.05, **p < 0.01, ***p < 0.001; black asterisks: FDR-corrected significance).

For place-selective ROIs, there was a weak trend for opposite hemispheric biases. Statistically, no differences between the groups were observed in MPA (p = 0.895) and PIGPA (p = 0.115). OPA showed a modest difference (p = 0.005; FDR-corrected p = 0.020), while PPA exhibited only a marginal effect (p = 0.050) that did not survive FDR correction (FDR-corrected p = 0.100) (Figure 6C). Thus, relative to the face-selective areas, evidence for differential hemispheric bias in place-selective areas was limited in both scope and magnitude. For body-selective ROIs, no differences were detected in either EBA (p = 0.336) or FBA (p = 0.795) (Figure 6D).

Together, these analyses demonstrate that hemispheric reorganization associated with atypical LN lateralization is selective rather than global. Robust and threshold-independent differences were consistently observed in the mSTS, with a secondary effect in the FFA, whereas other face-selective areas showed weaker, task-dependent, or no differences. The place– and body-selective areas also exhibited little to no evidence of systematic hemispheric bias differences between LHLD and RHLD participants.

## Discussion

The present study examined whether atypical lateralization of the language network (LN) is associated with global reorganization across other lateralized functions or instead gives rise to selective, region-specific changes. Recent work has argued that atypical language dominance is accompanied by global shifts in hemispheric organization at the level of macroscale functional gradients and large-scale cortical architecture (Labache et al., 2023). In contrast, using Human Connectome Project (HCP) data and converging analyses across tasks, thresholds, and levels of description, we found that LN-associated reorganization in category-selective visual network was highly selective rather than global. Hemispheric bias reversals were robust and consistent in the middle superior temporal sulcus (mSTS), accompanied by a secondary effect in the fusiform face area (FFA), whereas other face-selective areas showed weaker or task-dependent effects. Moreover, place– and body-selective areas showed little to no systematic modulation by LN lateralization. Taken together, these findings argue against a wholesale hemispheric inversion across category-selective visual areas and instead support a model in which atypical LN lateralization selectively reshapes cortical organization in regions that are anatomically and functionally positioned at the interface between LN and face-processing network.

### Selective reorganization within the face-processing network

Among all face-selective areas examined, the mSTS emerged as the locus of the most reliable and consistent hemispheric reorganization. The mSTS showed a clear reversal of hemispheric bias between left-hemisphere language dominance (LHLD) and right-hemisphere language dominance (RHLD) participants across both the working memory and emotion processing tasks. Importantly, the relationship between LN and face-processing network lateralization in the mSTS was not limited to categorical group differences, but instead followed a graded, continuous pattern across individuals, with stronger LH language dominance associated with stronger RH face-processing network bias, and vice versa. This convergence of group-level and correlational evidence indicates that the mSTS effect reflects a systematic organizational relationship rather than an artifact of group definition or threshold choice.

In contrast, the FFA showed a significant difference in the working memory task and a reliable, though weaker, association with LN lateralization across participants. aSTS, pSTS, MFA, and OFA showed little to no evidence of systematic hemispheric reorganization. This graded pattern across face-selective areas suggests that LN-associated reorganization does not uniformly propagate across the entire face-processing network, but instead preferentially affects a subset of regions with specific functional and anatomical properties.

### Spatial proximity and overlap with the language network

A key feature distinguishing the mSTS from other face-selective areas is its close spatial proximity to, and partial overlap with, core language-responsive network. Prior work has shown that regions within the posterior and middle portions of the superior temporal sulcus are engaged both by linguistic stimuli and by socially relevant visual information, including dynamic facial cues (Deen et al., 2015; Venezia et al., 2017). The present results extend this literature by demonstrating that this anatomical convergence has functional consequences for hemispheric organization: when the LN is lateralized to the RH, the mSTS shows a corresponding shift in face-processing bias toward the left hemisphere.

This spatial arrangement positions the mSTS at a functional intersection between LN and face-processing network. Regions located at such intersections may be especially susceptible to competitive or redistributive pressures during development or evolution, as they are recruited by multiple high-level cognitive domains. In this context, atypical lateralization of language may alter the balance of hemispheric specialization in adjacent or overlapping cortex, resulting in selective reorganization rather than global inversion.

### Competition for common neural resources

The observed pattern of hemispheric reorganization is consistent with theoretical frameworks emphasizing competition for shared neural resources. Models of biased competition propose that cortical representations vie for limited processing capacity, such that increased engagement or specialization of one representation can bias, suppress, or displace others, particularly within overlapping or closely adjacent cortical territories (Desimone and Duncan, 1995; Kastner and Ungerleider, 2001; Scalf et al., 2013). Within this framework, strong lateralization of LN to one hemisphere may bias face-processing network toward the contralateral hemisphere in regions where representational demands overlap or compete. A similar competitive process has also been reported between language and social processing in the brain (Rajimehr et al., 2022). Subjects with stronger language activations in some areas in LH tended to have stronger social activations in the homotopic areas in RH, leading to an opposite hemispheric lateralization for language and social processing.

Importantly, this account does not imply that LN and face-processing network share identical neural computations, but rather that their cortical implementations may draw on partially overlapping or neighboring neural populations. As illustrated in Figure 7, the mSTS face-selective area lies in close spatial proximity to LN, positioning it as a potential bottleneck where such competitive interactions are most likely to arise. The graded relationship observed between the language laterality index (LLI) and face laterality index (FLI) further supports this view, as it suggests a continuous trade-off rather than an all-or-none reallocation of function.

**Figure 7.**
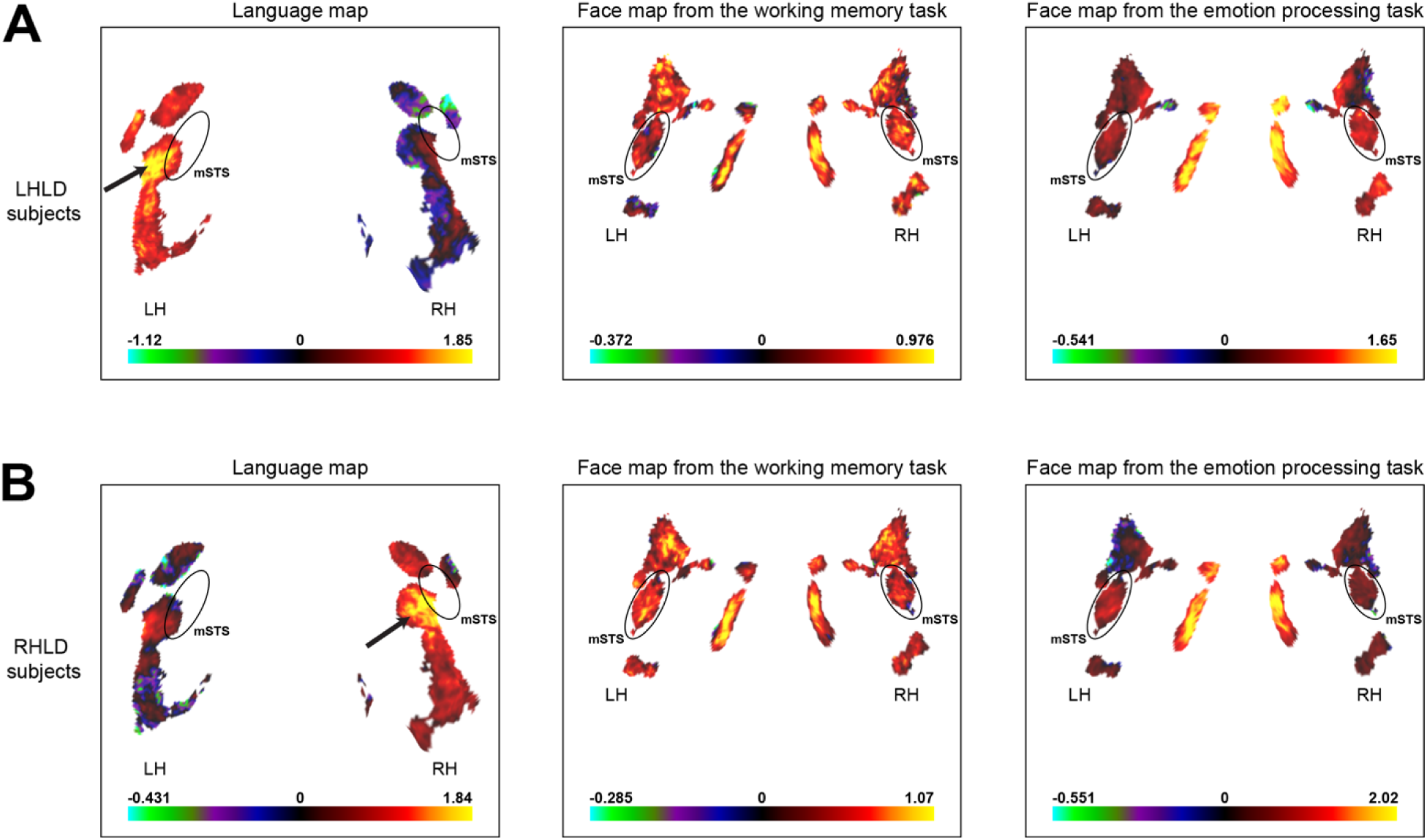
Topographic relationship between language and face activations in STS. The flat patches were cropped and magnified to show language activations, face activations from the working memory task, and face activations from the emotion processing task in LHLD participants (A) and RHLD participants (B). The cropped region highlights areas in lateral and ventral temporal cortex. The hotspot of language activation in STS is located adjacent to the face-selective mSTS.

### Secondary involvement of the fusiform face area

The involvement of the FFA, most clearly observed in the working memory task, likely reflects its strong functional and structural coupling with the mSTS rather than direct competition with the LN. Previous studies have consistently demonstrated structural connectivity and coordinated activity between the FFA and temporal face-selective areas during face perception and memory tasks, indicating that these regions operate as components of an integrated face-processing network (Blank et al., 2011; Turk-Browne et al., 2010; Wang et al., 2016).

Another line of evidence for the link between the mSTS and FFA comes from cross-species comparisons. It has been suggested that the human mSTS and FFA are homologues of middle temporal face patches in monkeys (Rajimehr et al., 2009; Yovel and Freiwald, 2013). These patches, including middle fundus (MF) and middle lateral (ML), are located adjacent to each other, are highly inter-connected, and complement each other in face processing (Hesse and Tsao, 2020). The human homologues of MF and ML—mSTS and FFA—have been spatially separated, presumably due to a disproportionate expansion of the intervening lateral temporal cortex in humans (Hill et al., 2010), yet they may have maintained strong structural and functional coupling over the course of evolution.

Within this framework, reorganization originating in the mSTS could influence the hemispheric bias of the FFA indirectly, particularly in tasks that place greater demands on integrative or memory-related aspects of face processing. The task dependence of the FFA effect observed here is consistent with this interpretation and suggests that coupling strength and task context modulate the extent to which reorganization propagates beyond the mSTS.

### Absence of global reorganization across category-selective visual network

Crucially, the present results provide little support for the hypothesis that atypical LN lateralization induces a global inversion of hemispheric specialization across category-selective visual network. Place-selective areas showed only weak and inconsistent differences, limited primarily to the occipital place area (OPA), and body-selective areas showed no reliable modulation by LN lateralization. These null or weak effects are informative: they indicate that LN-associated reorganization does not indiscriminately reshape hemispheric biases across all category-selective visual areas, even when those systems are themselves typically right-lateralized.

This selectivity argues against models positing a single, domain-general mechanism governing hemispheric specialization across the cortex. Instead, the data favor a more intricate view in which hemispheric organization emerges from local interactions among functionally and anatomically constrained systems. Under this view, atypical lateralization in one domain influences other domains primarily where there is close anatomical proximity, shared functional demands, or strong network-level coupling.

### Implications for theories of hemispheric specialization

Taken together, these findings refine our understanding of hemispheric specialization by demonstrating that atypical LN lateralization is associated with selective, region-specific reorganization rather than wholesale hemispheric reversal. The results highlight the importance of considering fine-grained cortical topology and network architecture when evaluating the consequences of atypical lateralization. Rather than treating hemispheric dominance as a global property of cerebral cortex, the present data suggest that lateralization is best understood as an emergent property of interacting networks, shaped by local constraints and competitive dynamics.

More broadly, this work underscores the value of studying atypical populations—not as deviations from a norm, but as a valuable group of individuals who reveal organizing principles of the human brain. By examining how LN and face-processing network interact under conditions of atypical lateralization, we gain insight into the flexibility and limits of cortical specialization.

## Methods

### Participants

Data were obtained from the Human Connectome Project (HCP) S1200 release. Participants with available task-fMRI data for the HCP Language task (story vs. baseline contrast) were included. The full sample used to characterize the distribution of language lateralization comprised N = 1,049 participants. The HCP data were acquired using protocols approved by the Washington University institutional review board, and written informed consent was obtained from all subjects.

For group-based analyses, participants exhibiting strong hemispheric dominance for language were identified based on the LLI. Individuals with LLI values more than two standard deviations above or below the sample mean were classified as having left-hemisphere language dominance (LHLD) or right-hemisphere language dominance (RHLD), respectively. This procedure identified 17 LHLD and 28 RHLD participants. To obtain matched group sizes, the LHLD group was supplemented with additional participants exhibiting the highest positive LLI values, yielding 28 LHLD and 28 RHLD participants for between-group comparisons.

Complete task-fMRI data for group-level analyses were available for 27 LHLD–RHLD participant pairs in the working memory task and 28 LHLD–RHLD participant pairs in the emotion processing task, and these samples were used for all corresponding analyses.

### Language task

Functional data in this study were based on the HCP language processing task (Binder et al., 2011). The language task consisted of two runs (run duration = 3:57 min:sec). In each run, 4 blocks of a story task were interleaved with 4 blocks of a math task. The lengths of blocks varied, and the average duration of blocks was approximately 30 s.

In the story blocks, participants were presented with brief auditory stories (5–9 sentences) adapted from Aesop’s fables, followed by a 2-alternative forced-choice question that asked participants about the topic of the story. For example, after a story about an eagle that saves a man who had done him a favor, participants were asked “Was that about revenge or reciprocity?” Participants pressed a button to select either the first or the second choice. The math task also included trials that were presented auditorily. In these trials, participants completed a series of simple arithmetic (addition and subtraction) operations (e.g., “Fourteen plus twelve”), followed by “equals” and then two choices (e.g., “twenty-nine or twenty-six”). Participants pressed a button to select either the first or the second answer. The math task was adaptive to maintain a similar level of difficulty across participants. The math condition served as a control that matched the story condition for auditory input, motor response demands, and block duration, while minimizing language-specific semantic processing.

### Working memory task

The HCP working memory task combined a category-selective visual localizer with a working-memory manipulation. Participants viewed blocks of images from four stimulus categories—faces, places, tools, and body parts—with categories presented in separate blocks within each run. Task demands alternated between a 2-back condition (working memory) and a 0-back condition (control). At the start of each block, a brief cue indicated the task condition and (for 0-back) the target stimulus. Each run consisted of task blocks interleaved with fixation periods. Within each trial, a stimulus was presented briefly, followed by a short inter-trial interval. Targets in the 2-back condition were defined as repeats with a two-trial lag, whereas in the 0-back condition targets matched the cued stimulus; non-targets were non-matching items. The lure events were included to increase decision demands (e.g., near-repeat sequences in 2-back or repeated non-target items in 0-back). For the present analyses, category-selective activation was quantified using the FACE-AVG, PLACE-AVG, and BODY-AVG contrasts, collapsing across memory conditions.

### Emotion processing task

The HCP Emotion Processing task was adapted from the paradigm developed by Hariri and colleagues and used a block design requiring perceptual matching (Hariri et al., 2006; Manuck et al., 2007). Participants completed blocks in which they matched either emotional faces (angry or fearful expressions) or geometric shapes. In each block, a cue indicated whether the upcoming trials involved face matching or shape matching. Trials presented a target stimulus and two choice stimuli, and participants selected the matching option. Face and shape blocks were alternated across two runs, with fixation epochs included at the end of each run. In the present study, face-related responses were assessed using the standard faces vs. shapes contrast.

### Data acquisition

Images were acquired using a customized 3T Siemens ‘Connectom’ Skyra scanner having a 100 mT/m SC72 gradient insert and a standard Siemens 32-channel RF-receive head coil. At least one 3D T1-weighted MPRAGE image and one 3D T2-weighted SPACE image were acquired at 0.7 mm isotropic resolution. Whole-brain task fMRI data were acquired using a multi-band EPI sequence with parameters of TR = 720 ms, TE = 33.1 ms, flip angle = 52°, 2 mm isotropic voxels, 72 slices, and multi-band acceleration factor of 8. Spin-echo field maps were acquired during both structural and fMRI scanning sessions to enable accurate cross-modal registration of structural and functional images in each subject.

### Analysis of structural data

Structural images (T1-weighted and T2-weighted) were used for extracting subcortical gray matter structures and reconstructing cortical surfaces in each subject. Volume data were transformed from native space into MNI space using a nonlinear volume-based registration. For accurate cross-subject registration of cortical surfaces, a multimodal surface matching (MSM) algorithm was used. The MSM algorithm had two versions: ‘MSMSulc’ (non-rigid surface alignment based on folding patterns) and ‘MSMAll’ (optimized alignment of cortical areas using sulcal depth maps plus features from other modalities including myelin maps, resting-state network maps, and visuotopic connectivity maps). Data in our work were based on MSMAll registration. After surface and volume registration, cortical vertices were combined with subcortical gray matter voxels to form the standard ‘CIFTI grayordinates’ space (91,282 vertices/voxels with ∼2 mm cortical vertex spacing and 2 mm isotropic subcortical voxels).

### Analysis of fMRI data

Functional images were minimally preprocessed using the HCP pipelines (Glasser et al., 2013). Preprocessing included correction for spatial distortions due to gradient nonlinearity and B0 field inhomogeneity, fieldmap-based unwarping of EPI images, motion correction, brain-boundary-based registration of EPI to structural T1-weighted scans, non-linear registration to MNI space, and grand-mean intensity normalization. Data from the cortical gray matter ribbon were projected onto the surface and then onto the standard grayordinates space. Data were minimally smoothed by a 2 mm FWHM Gaussian kernel in the grayordinates space. Thus, smoothing was constrained to the cortical surface mesh in each hemisphere.

The preprocessed functional time-series were entered into a general linear model (GLM) to estimate functional activities in each vertex/voxel in each run. Two regressors were included in the GLM design of the language task: story and math. For the working memory task, eight regressors were used in the GLM design—one for each type of stimulus in each of the N-back conditions. For the emotion processing task, two regressors were used in the GLM design: emotional faces and shapes. Each regressor covered the duration of a block. All regressors were convolved with a canonical hemodynamic response function and its temporal derivatives. The time-series were temporally filtered with a Gaussian-weighted linear high-pass filter with a cutoff of 200 s, to remove low-frequency drifts/fluctuations presumably unrelated to the task design. The time-series were also prewhitened to remove temporal autocorrelations in the fMRI signal.

## Author’s contributions

Parham Zargar conceived the idea, performed the analyses, and wrote the manuscript. Reza Rajimehr designed the study, supervised the project, and critically revised the manuscript.

## Data availability statement

This study used publicly available data from the Human Connectome Project (HCP). Access to the HCP dataset can be obtained through the official HCP data portal. The analysis codes developed for this study are available upon reasonable request.

## Acknowledgments

We would like to thank Arsalan Firoozi, Roza Hamidi, and Mohammad Ebrahim Katebi for their assistance with data analysis and for their helpful comments. This research was supported by the Institute for Research in Fundamental Sciences (IPM).

